# A Functionally Conserved Mechanism of Modulation via a Vestibule Site in Pentameric Ligand-Gated Ion Channels

**DOI:** 10.1101/770719

**Authors:** Marijke Brams, Cedric Govaerts, Kumiko Kambara, Kerry Price, Radovan Spurny, Anant Gharpure, Els Pardon, Genevieve L. Evans, Daniel Bertrand, Sarah C. R. Lummis, Ryan E. Hibbs, Jan Steyaert, Chris Ulens

## Abstract

Pentameric ligand-gated ion channels (pLGICs) belong to a class of ion channels involved in fast synaptic signaling in the central and peripheral nervous systems. Molecules acting as allosteric modulators target binding sites that are remote from the neurotransmitter binding site, but functionally affect coupling of ligand binding to channel opening. Here, we investigated an allosteric binding site in the ion channel vestibule, which has converged from a series of studies on prokaryote and eukaryote channel homologs. We discovered single domain antibodies, called nanobodies, which are functionally active as allosteric modulators, and solved co-crystal structures of the prokaryote channel ELIC bound either to a positive (PAM) or a negative (NAM) allosteric modulator. We extrapolate the functional importance of the vestibule binding site to eukaryote ion channels, suggesting a conserved mechanism of allosteric modulation. This work identifies key elements of allosteric binding sites and extends drug design possibilities in pLGICs using nanobodies.

## INTRODUCTION

In 1965, Monod, Wyman and Changeux postulated the model of allosteric modulation in proteins (Monod, Wyman, and Changeux 1965). According to this model, proteins exist in two possible conformational states, the tensed (T) or relaxed state (R). The substrate typically has a high affinity for the T state. In multi-subunit proteins, all subunits undergo a concerted transition from the R to the T state upon substrate binding. The equilibrium can be shifted to the R or the T state through a ligand that binds at a site that is different from the substrate binding site, in other words an allosteric site.

Changeux subsequently devoted much of his scientific career to the study of allosteric proteins, with specific attention begin paid to the nicotinic acetylcholine receptor (nAChR). This protein is a ligand-gated ion channel (LGIC) and thus in effect has no substrate, but the principle of allosteric modulation is similar in that binding of acetylcholine (ACh) shifts the thermodynamic equilibrium from a closed channel state to an open channel state through binding to a site ∼50 Å from the channel gate. The nAChR is a member of a superfamily of pentameric LGICs (pLGICs), which play major roles in fast synaptic transmission in the central and peripheral nervous systems, and are the site of action of many therapeutic drugs.

Structures of these proteins have been elucidated over the last decade and several nAChR structures are now available for the heteromeric α4β2 nAChR (Morales-Perez, Noviello, and Hibbs 2016; Walsh et al. 2018), as well as other members of the pLGIC family, including the 5-HT_3_ serotonin receptor (Hassaine et al. 2014; Polovinkin et al. 2018; Basak et al. 2018), the glycine receptor (Du et al. 2015; Huang et al. 2015), the GABA_A_ receptor (Zhu et al. 2018; Phulera et al. 2018; Laverty et al. 2019; Masiulis et al. 2019; Miller and Aricescu 2014), the glutamate-gated chloride channel from *C. elegans* GluCl (Hibbs and Gouaux 2011; Althoff et al. 2014) and the prokaryote channels ELIC (Hilf and Dutzler 2008) and GLIC (Hilf and Dutzler 2009; Bocquet et al. 2009). Historically, crucial structural insight into the class of nicotinic receptors was derived from cryo-electron microscopic structures of the nAChRs from the electric organ of *Torpedo* (Miyazawa, Fujiyoshi, and Unwin 2003; Unwin 2005) and X-ray crystal structures of the acetylcholine binding protein (AChBP; found in certain snails and worms), which is homologous to the extracellular ligand binding domain (LBD) of nAChRs (Brejc et al. 2001).

The concept of allosteric modulation is now also more broadly applied to understand the mode of action of certain drugs, called allosteric modulators, which bind at a site that is different from the neurotransmitter binding site, but which can alter energy barriers between multiple conformational states (Bertrand and Gopalakrishnan 2007). For example, in the case of pLGICs, positive allosteric modulators (PAMs) of the nAChR can facilitate a transition from a resting to an activated state, thus enhancing the agonist-evoked response. In contrast, negative allosteric modulators (NAMs) hinder such a transition, thus diminishing the agonist response. From a drug development perspective, PAMs or NAMs are highly attractive as they finely tune receptor activation without affecting the normal fluctuations of neurotransmitter release at the synapse. One of the most extensively described PAMs used in clinic are the benzodiazepines, which act on GABA_A_ receptors and are widely prescribed as hypnotics, anxiolytics, anti-epileptics or muscle relaxants. Important insights into the molecular recognition of these modulators are now revealed by high resolution structural data (Zhu et al. 2018; Phulera et al. 2018; Laverty et al. 2019; Masiulis et al. 2019; Miller and Aricescu 2014).

With the availability of a growing amount of structural data for these receptors, a diverse array of molecules has been revealed, many of which bind at distinct allosteric binding sites, including general anesthetics (Nury et al. 2011; Sauguet et al. 2013; Spurny et al. 2013), neurosteroids (Miller et al. 2017; Laverty et al. 2017; Q. Chen et al. 2018), lipids (Zhu et al. 2018; Laverty et al. 2019), antiparasitics (Hibbs and Gouaux 2011; Althoff et al. 2014), and many others (Nys et al. 2016). Detailed investigation of allosteric sites not only brings further knowledge about the receptor functionality but also uncovers novel drug target sites. However, our current understanding of this multi-site mechanism of allosteric modulation in pLGICs is still incomplete.

In this study, we used complementary structural and functional approaches to expand our understanding of the molecular mechanism of allosteric modulation in pLGICs. Using the prokaryote ELIC channel as a model, we explored the potential of nanobodies (single chain antibodies) as allosteric modulators. We discovered functionally active nanobodies, which act either as a PAM or NAM on ELIC and determined co-crystal structures to elucidate the nanobody interactions with ELIC. One of the structures reveals an allosteric binding site located near the vestibule of the extracellular ligand-binding domain. Comparison of conservation and divergence in this site in different prokaryotic and eukaryotic receptors suggests a mechanism for achieving subtype-selective allosteric modulation across the receptor superfamily. Using cysteine-scanning mutagenesis and electrophysiological recordings we extrapolate to how the vestibule site can also be targeted for modulation of the human 5-HT_3A_ receptor as a proof of principle relevant to other eukaryotic receptors.

## RESULTS AND DISCUSSION

### Identification of nanobodies active as allosteric modulators on ELIC

In this study, we took advantage of nanobodies, which are high affinity single chain antibodies derived from llamas; they have been widely employed to facilitate structural studies (Manglik, Kobilka, and Steyaert 2017) and also hold potential as therapeutics against many possible targets. A first example, caplacizumab (Cabilivi^®^), has recently reached the market (Scully et al. 2019). Using the prokaryote ion channel ELIC as a model system, we investigated whether nanobodies could be selected with allosteric modulator activity on ligand-gated ion channels. We expressed ELIC channels in *Xenopus* oocytes and employed automated electrophysiological recordings to characterize a panel of more than 20 different ELIC nanobodies. While none of the nanobodies had any functional effect on ELIC when applied alone, we found that co-application with the agonist GABA evoked a response that broadly falls into 3 categories. One type of nanobodies enhanced the agonist-evoked response, while a 2^nd^ type of nanobodies diminished the agonist-evoked response and the 3^rd^ type had little to no effect. From these, we selected several enhancers (PAMs) and inhibitors (NAMs) for a detailed electrophysiological characterization as potential allosteric modulators. In parallel, we conducted X-ray diffraction screening of ELIC plus nanobody co-crystals for structural elucidation. From this selection, we obtained a PAM-active nanobody (PAM-Nb) as well as another NAM-active nanobody (NAM-Nb) and determined their structures bound to ELIC by X-ray crystallography.

Co-application of the agonist GABA with a range of PAM-Nb concentrations demonstrates that PAM-Nb enhances the agonist response (Fig. 1a) with a pEC_50_-value of 5.37±0.03 (EC_50_: 4.2 µM) and I_max_=257±14% (n=8). In contrast, co-application of a range of NAM-Nb concentrations demonstrates that NAM-Nb decreases the agonist response (Fig. 1b) with a pIC_50_-value of 6.89±0.03 (IC_50_: 0.13 µM) and I_max_=34±2%% (n=6). Unlike competitive antagonists, which fully inhibit the agonist response at saturating concentrations, the inhibition of NAM-Nb levels off at 70% of the response with GABA alone, consistent with the mode of action of certain NAMs. These results demonstrate that functionally active nanobodies can be developed against the ligand-gated ion channel ELIC.

**Figure 1.**
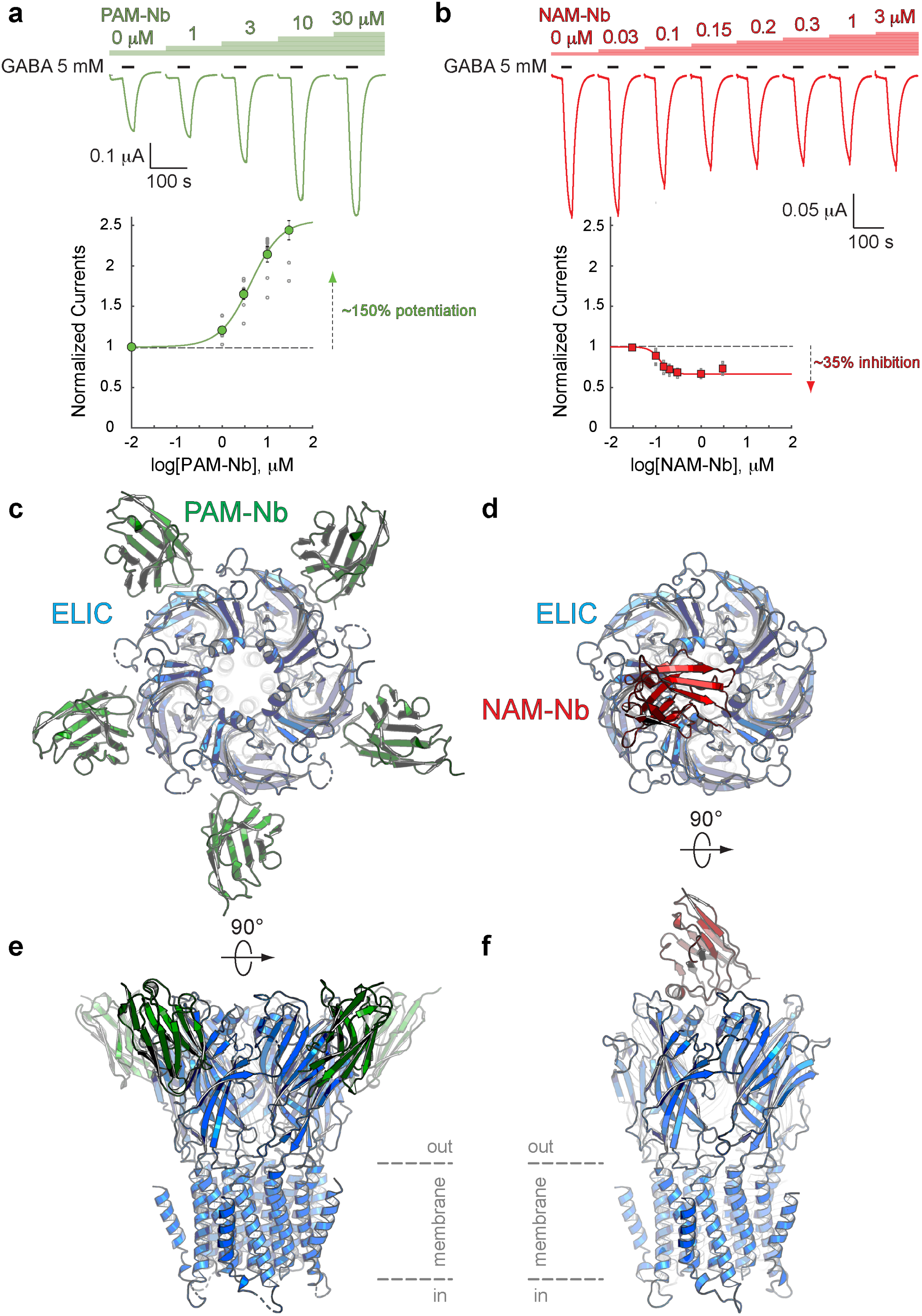
Nanobodies active as allosteric modulators and structures bound to the ELIC channel. **a, b.** Electrophysiological recordings of ELIC activated by the agonist GABA and in the presence of increasing concentrations of PAM-Nb (a, green) or NAM-Nb (b, red). The curve represents a fit to the Hill equation to the normalized current responses. Circles represent averaged data with standard errors. **c,d.** X-ray crystal structures of ELIC bound by PAM-Nb (c) or NAM-Nb (d). The cartoon representation shows a top-down view onto the ELIC pentamer along the 5-fold axis (blue). **e,f.** Side views from **c,d**. The dashed lines indicate the presumed location of the membrane boundaries.

### Crystal structures of ELIC in complex with a PAM- or NAM-type nanobody

To gain insight into the structural recognition of positive (PAM-Nb) or negative (NAM-Nb) allosteric modulators we solved X-ray crystal structures of ELIC in complex with the PAM-Nb or NAM-Nb, respectively. The structure of ELIC in complex with the PAM-Nb was determined at a resolution of 2.59Å, and reveals five PAM-Nb molecules bound to a single pentameric ELIC channel (Fig. 1c,e). Each PAM-Nb binds at an intrasubunit site in the ELIC extracellular ligand binding domain. When viewed from the top along the 5-fold symmetry axis the nanobodies extend outward and the structure resembles a 5-bladed propeller (Fig. 1c). The structure of ELIC in complex with NAM-Nb was determined at a resolution of 3.25Å and is structurally distinct from the complex with PAM-Nb: instead of five Nb molecules bound to the ELIC pentamer, here a single NAM-Nb molecule is bound to the ELIC pentamer (Fig. 1d,f). The NAM-Nb binds at the channel vestibule entrance and near to the N-terminal α-helix of a single ELIC subunit, and is oriented in such a manner that only a single nanobody molecule can bind at this interface, as the core of the nanobody sterically hinders access to the 4 other sites (Fig. 1d,f).

A more detailed analysis of the interaction interface between both nanobodies and ELIC reveals remarkable features (Fig. 2). The PAM-Nb binds to the extracellular ligand-binding domain and forms extensive interactions through the complementarity determining region CDR1 (residues S25-I33) of the nanobody (Fig. 2a). The tip of the finger of this loop region wedges in between the ELIC β8- and β9-strand, forming a distinct anti-parallel β-sheet interaction with the β8-strand. The CDR1 loop region points toward an allosteric binding site previously identified in a chimera of the human α7 nAChR and the acetylcholine binding protein, α7-AChBP (Li et al. 2011) (see complex with fragment molecule CU2017, pdb code 5oui) (Fig. 2a). In other words, the PAM-Nb binds to a site in ELIC that corresponds to a functionally important allosteric site in the human α7 nAChR, consistent with its function as an allosteric modulator.

**Figure 2.**
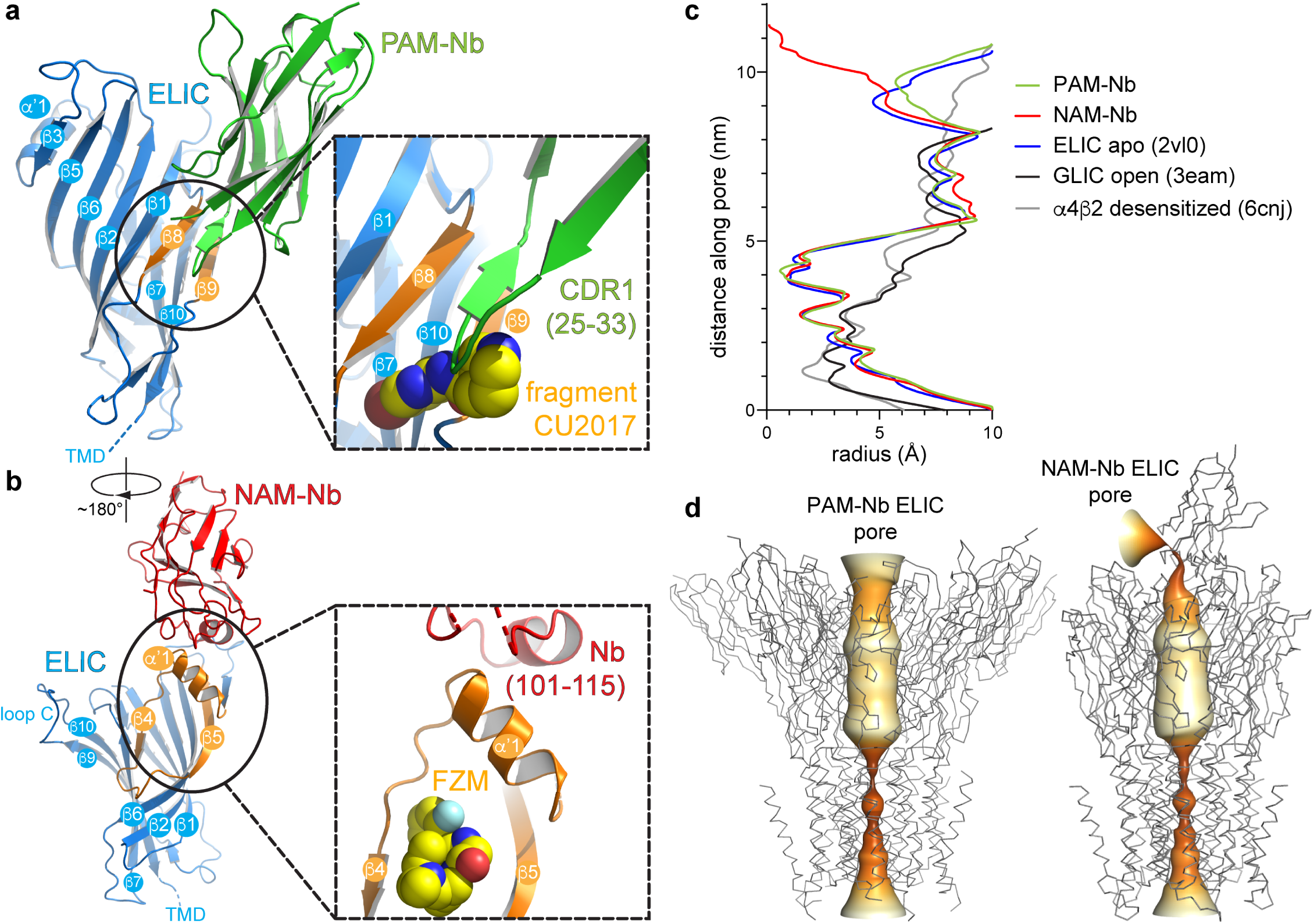
Detailed nanobody interaction sites in ELIC and channel pore analysis. **a.** A detailed view of the interaction between PAM-Nb (green) and a single ELIC subunit (blue). The binding site for PAM-Nb overlaps with a known allosteric binding site for a small molecule fragment called CU2017 (ref. (Delbart et al. 2018), pdb code 5oui), bound between the β8-β9 strands (in orange) and the fragment shown as spheres (carbon yellow, nitrogen blue). **b.** Detailed view of the interaction between NAM-Nb (red) and a single ELIC subunit (blue). The binding site for NAM-Nb involves a region which is part of a known allosteric binding site for flurazepam (ref. (Spurny et al. 2012), pdb code 2yoe). **c,d.** Analysis of the ELIC channel pore radius for PAM-Nb, NAM-Nb bound structures and references structures.

The mode of interaction of the NAM-Nb with ELIC is distinct from the PAM-Nb (Fig. 2b). The interaction interface is remarkable in that none of the CDR1-, CDR2- or CDR3-regions interact with ELIC. Instead, it is the region P104-L110 of the NAM-Nb that forms an α-helix, and which corresponds to a region that has no defined secondary structure in most other nanobody structures. This α-helical region interacts with the α’1-helix in ELIC (N60-N69) and forms the outer rim of an allosteric binding site previously identified as the vestibule binding site (Spurny et al. 2012). This site is the target for the benzodiazepine flurazepam, which acts as a positive allosteric modulator on ELIC (Spurny et al. 2012) (Fig. 2b). Similar to the PAM-Nb, it is interesting to observe that the allosteric binding site of the NAM-Nb also overlaps with a previously identified binding site for a small molecule allosteric modulator. Remarkably, in each case, the nanobody has the opposite functional effect of the small molecule at the same site. The PAM-Nb acts as a positive modulator, whereas the CU2017 fragment acts as a negative modulator at the β8-β9 site. Conversely, the NAM-Nb acts as a negative modulator, whereas flurazepam acts as a positive modulator at the vestibule site. This result demonstrates that the same allosteric site can be targeted both by a PAM or NAM, and that its functional action likely depends on defined side chain interactions. This is consistent with previous pharmacological studies, which have shown that a substitution as small as a methylation of an aromatic ring in a small molecule modulator can alter the functional profile from a PAM to a NAM of the α7 nAChR (Gill-Thind et al. 2015).

The pores of both Nb complexes resemble previous structures of ELIC, with narrow constriction points at the 9’, 16’, and 20’ positions (Hilf and Dutzler 2008), suggesting that the channels are in non-conducting conformations (Fig. 2c,d). This closed state is structurally distinct from the putative desensitized state of the α4β2 nAChR, whose primary gate lies at the cytoplasmic end at the −1’ position^3^. Further analysis of the central channel axis of the NAM-Nb complex reveals a complete occlusion of the extracellular vestibule by the nanobody (Fig. 2d). Despite this apparent block of the ion permeation pathway, NAM-Nb only inhibits GABA-induced currents by ∼35%, as mentioned previously. This can potentially be explained by alternate pathways for ion entry through lateral fenestrations located at subunit interfaces, as seen in other members of the pLGIC family (Zhu et al. 2018; Miller and Aricescu 2014).

### Subtype-dependent vestibule site access in different prokaryote and eukaryote receptors

To further investigate the possible conservation of the vestibule binding site in different prokaryote and eukaryote pLGICs, we performed a systematic analysis of the vestibule site architecture in all currently available pLGIC structures. The results from this analysis show that the outer rim of the vestibule site, which corresponds to residues N60-F95 in ELIC, can adopt one of 3 possible conformations (Fig. 3).

**Figure 3.**
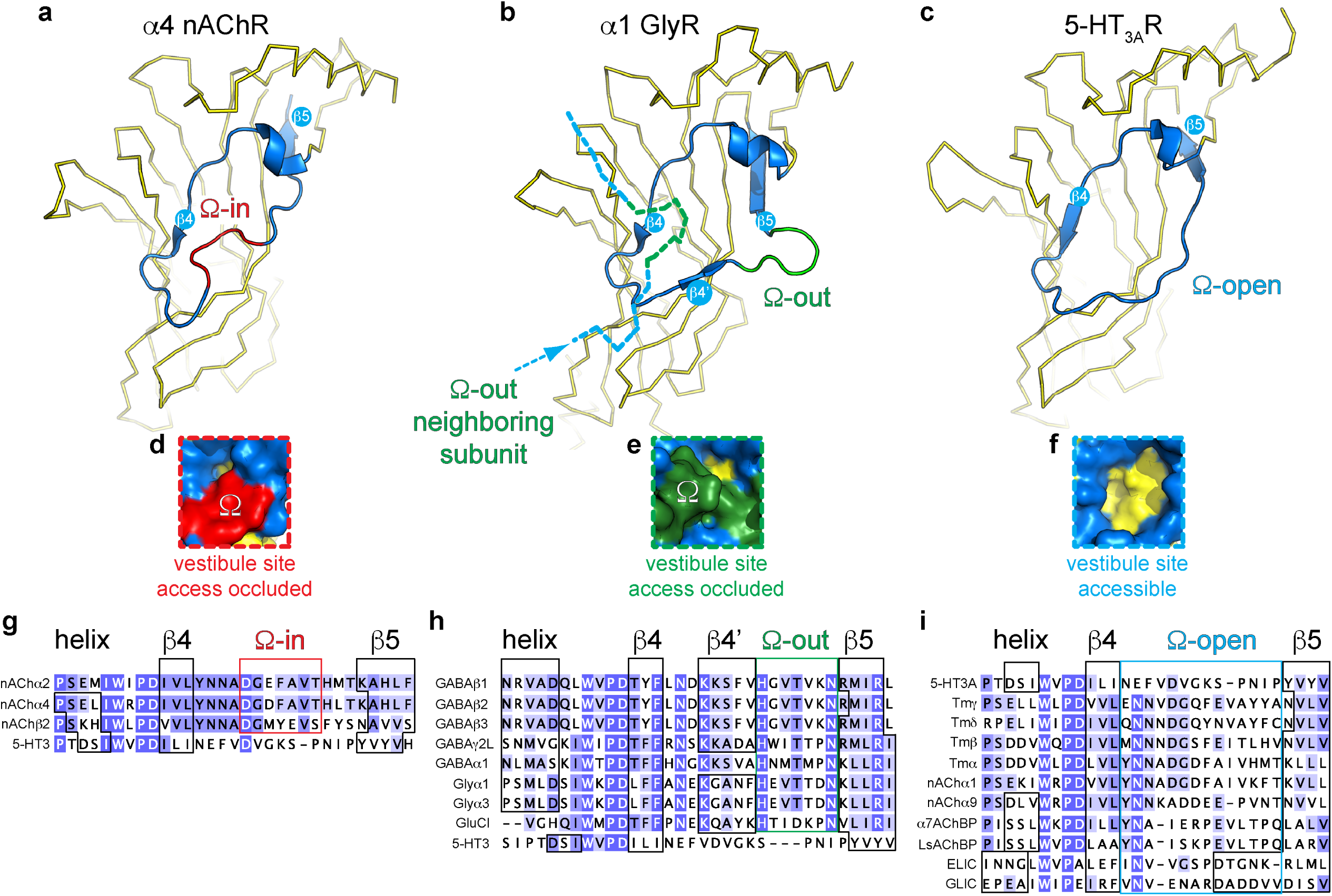
Distinct conformations of the vestibule site in pentameric ligand-gated ion channels. **a-c.** Yellow ribbon representation of a single subunit ligand binding domain. Part of the vestibule site (shown in blue cartoon), called the Ω-loop, adopts 3 distinct conformations in different pLGICs: the Ω-in (red, **a**), Ω-out (green, **b**) and Ω-open conformation (blue, **c**). **d-f**. Insets show a zoom of the Ω-loop in surface representation to illustrate occluded vestibule site access in the Ω-in and Ω-out conformations, compared to an accessible vestibule site in the Ω-open conformation. **g-i**. Sequence alignment of the Ω-loop and neighboring residues in pLGICs for which structures have been elucidated.

In certain structures, we observe that the stretch of amino acids between the β4- and β5-strands resembles the shape of the Greek letter Ω, for example in the α4 nAChR subunit (Morales-Perez, Noviello, and Hibbs 2016; Walsh et al. 2018), and therefore this region is termed the Ω loop (Hu et al. 2018). In Fig. 3a, the outer rim of the vestibule site in the α4 nAChR is shown in blue (from the helix to the β5-strand) and the Ω loop in red. The tip of the Ω loop can either point into the vestibule as in the α4 nAChR subunit (Morales-Perez, Noviello, and Hibbs 2016; Walsh et al. 2018), which we call the Ω-in conformation (Fig. 3a), or the tip of the Ω loop can point outward, which we call the Ω-out conformation, for example in the α1 GlyR subunit (Du et al. 2015) (outer rim shown in blue, Ω-out in green, Fig. 3b). Importantly, in both of these conformations access to the vestibule site is occluded. In the Ω-in conformation, the tip of the Ω loop prevents access to the vestibule site of its own α4 nAChR-subunit (Fig. 3a,d), whereas in the Ω-out conformation vestibule access is prevented by the tip of the Ω-out loop of its neighboring α1 GlyR (-) subunit (Fig. 3b,e). In addition to the Ω-in and Ω-out conformations, we observe a third possible conformation in which the Ω loop is stretched, for example in the 5-HT_3A_R (Hassaine et al. 2014) (Fig. 3c) and ELIC, and creates an accessible vestibule site (Fig. 3f). Consequently, we term this conformation the Ω-open conformation (outer rim and Ω-open is shown in blue, Fig. 3c,f).

A systematic comparison of the Ω loop conformation demonstrates that the different pLGIC subfamilies group into defined categories. First, we observe that all currently known anionic receptor structures adopt the Ω-out conformation (Fig. 3h). This implies that the vestibule access is occluded in all of these receptors, for example in the homomeric α1 GlyR (Du et al. 2015), α3 GlyR (Huang et al. 2015), or GluCl (Hibbs and Gouaux 2011) as well as the heteropentameric α1βγ2 GABA_A_Rs (Zhu et al. 2018; Phulera et al. 2018; Laverty et al. 2019) and homopentameric β3 GABA_A_R (Miller and Aricescu 2014). The Ω-out loop sequence is strongly conserved in these receptors, with a His residue and Asn residue at either end of the Ω loop and a Thr at the tip (not conserved in GluCl). We also observe that in all these cases the Ω loop is preceded by a stretch of amino acids that forms an additional β-sheet, which we call the β4’-sheet, as it also follows the β4-sheet. The β4’-sheet contains a well conserved start Lys residue, which is present in GlyRs, GABA_A_Rs and GluCl. The sequence conservation in these 2 regions suggests an important functional role. In contrast with anionic receptors, we observe that certain cationic receptors can adopt either the Ω- in or Ω-open conformation. However, none of the cationic receptors adopt the Ω-out conformation. Only 3 nAChRs adopt the Ω-in conformation, namely the α2 nAChR (Kouvatsos et al. 2016), α4 and β2 nAChR-subunits (Morales-Perez, Noviello, and Hibbs 2016; Walsh et al. 2018). This implies that the heteropentameric α4β2 nAChRs (Morales-Perez, Noviello, and Hibbs 2016; Walsh et al. 2018) adopt an all-subunit-occluded vestibule site conformation. The stretch of amino acids that form the Ω-in conformation is also well conserved with an Asp-Gly and Val-Thr/Ser on either end and an aromatic residue (Phe/Tyr) at the tip, again suggesting an important functional role. In contrast, all other nAChR subunits (Dellisanti et al. 2007; Zouridakis et al. 2014), including the Torpedo nAChR (Miyazawa, Fujiyoshi, and Unwin 2003), as well as the 5-HT_3A_R (Hassaine et al. 2014), snail AChBPs (Brejc et al. 2001; Celie et al. 2005), prokaryote GLIC (Bocquet et al. 2009; Hilf and Dutzler 2009) and ELIC adopt the Ω-open conformation. The sequence of amino acids forming the Ω-open loop is less conserved, except for the position +2 following the β4-sheet, which is a Asn in all receptors except 5-HT_3A_R. Important to note is that the Ω-open loop in the α9 nAChR (Zouridakis et al. 2014) is disordered in the apo state (pdb code 4d01), suggesting flexibility in this region, but not in the antagonist (MLA)-bound state (pdb code 4uxu). This raises the intriguing possibility that the Ω loop is intrinsically flexible and could undergo drug-induced conformational changes, similar to loop C at the neurotransmitter binding site (Brams et al. 2011).

Together, the results from this analysis demonstrate that different receptor subtypes adopt a different Ω loop conformation, with the Ω-open conformation creating an accessible vestibule site, whereas the Ω-in and Ω-out conformations occlude vestibule site access. This discovery provides new opportunities for drug design of allosteric molecules that have subtype-specific pharmacology (due to low sequence conservation compared to the orthosteric site), based on vestibule site access.

### Cysteine-scanning mutagenesis in the vestibule site of the human 5-HT_3A_ receptor

Based on our observations that the vestibule site in ELIC is the target for positive allosteric modulators such as the benzodiazepine flurazepam (Spurny et al. 2012) as well as negative allosteric modulators such as the NAM-nanobody described in this study, we further investigated whether the mechanism of vestibule site modulation is functionally conserved in eukaryote receptors. We chose the 5-HT_3A_R as a prototype receptor since it has a clear Ω-open vestibule conformation and its structure (Hassaine et al. 2014) as well as functional and pharmacological properties (Lummis 2012) are well described. We then employed the substituted cysteine accessibility method (SCAM) (Karlin and Akabas 1998) to investigate the functional effects on channel gating before and after modification of cysteines in the 5-HT_3A_R vestibule site with the cysteine-reactive reagent MTSEA-biotin. We chose residues on the outer rim of the vestibule site, T112 (top) and F125 (bottom), respectively, as well as residues deeper into the vestibule site, N147, K149 and L151 on the β6-strand and Y86 on the β2-strand (Fig. 4b). It is interesting to note that N147, K149 and L151 are located on the opposite side of the β-strand to the loop E residues (Q146 and Y148), while Y86 is on the opposite side of the β-strand to the loop D residues (W85 and R87); both of these regions are functionally important contributors to the neurotransmitter binding site (Hassaine et al. 2014). Residue P111, which points away from the vestibule site, was included as a negative control.

**Figure 4.**
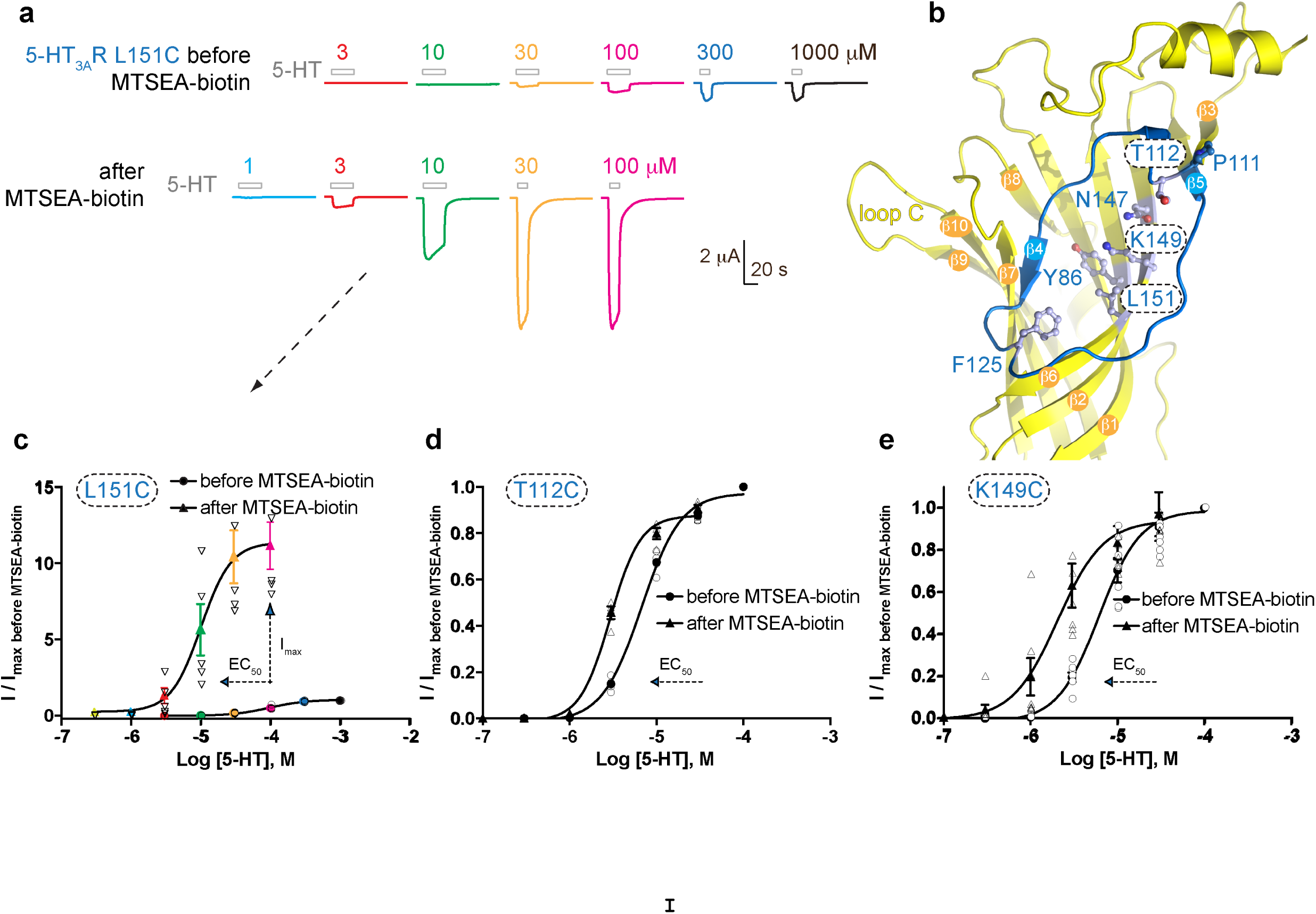
Allosteric modulation of the 5-HT_3_A receptor through chemical modification of engineered cysteines in the vestibule site. **a.** Example traces of agonist-evoked channel responses of the L151C 5-HT_3A_R mutant before and after modification with MTSEA-biotin show potentiation after cysteine modification. **b**. Location of L151C and other engineered cysteine mutants in this study, shown in ball&stick representation. **c**. Concentration-activation curves before and after modification with MTSEA-biotin are a Hill curve fit for the recordings shown in (**a**) as well as additional data for T112C (**d**) and K149C. (**c-e**) Each of these 3 mutants reveal a leftward shift of the curve upon cysteine modification, consistent with a positive allosteric effect. In the case of L151C this effect is combined with a large increase of the maximal current response.

Representative current traces of channel responses to increasing concentrations of serotonin (5-HT) are shown in Fig. 4a for the L151C mutant before and after modification with MTSEA-biotin. The channel responses are drastically increased after cysteine modification, which is caused by a more than 10-fold increase of the maximal current response (I_max_) of the concentration-activation curve after MTSEA-biotin modification as well as a shift of the EC_50_-value to lower concentrations (Fig. 4a,c) (pEC_50_: 4.01 ± 0.03, n=5 (EC_50_: 98 µM) versus pEC_50_: 4.96 ± 0.04, n= 7 (EC_50_: 11 µM)), indicating a strong PAM-effect at this position (*P* < 0.001). Mutants T112C and K149C also showed a significant decrease of the EC_50_-value after MTSEA-biotin, namely 6.7 µM (pEC_50_: 5.17 ± 0.02, n=4) versus 2.9 µM (pEC_50_: 5.53 ± 0.02, n=4) for T112C (Fig. 3d) (*P* < 0.0001) and 6.2 µM (pEC_50_: 5.21 ± 0.05, n=7) versus 2.1 µM (pEC_50_: 5.69 ± 0.11, n=7) for K149C (Fig. 3e) (*P* < 0.002). Although neither of these mutants displayed an increase of I_max_ as in the L151C mutant, the leftward shift of the concentration-activation curve is also consistent with a PAM-effect at these positions. No significant differences were observed for current responses before and after MTSEA-biotin for Y86C, F125C and N147C or the negative control, P111C, suggesting modification of these residues does not place the MTSEA-biotin in an appropriate position to act as a modulator, although it is also possible it did not react here. Together, these results demonstrate that allosteric modulation via the vestibule binding site is functionally conserved between ELIC and 5-HT_3A_ receptors. The evolutionary distance between these receptors, combined with previous work, suggests that this mechanism of modulation likely also extends to other pLGIC members, and indeed the functional importance of the vestibule site in channel modulation is now emerging from a wide range of different studies demonstrating ELIC modulation by flurazepam (Spurny et al. 2012) or nanobodies (this study), 4-bromocinnamate modulation of the prokaryote homolog sTeLIC (Hu et al. 2018), as well as eukaryote 5-HT_3A_R modulation by cysteine-scanning mutagenesis (this study), and α7 nAChR modulation using a fragment-based screening approach (Spurny et al. 2015; Delbart et al. 2018).

In conclusion, we demonstrate here that functionally active nanobodies can be developed for the ligand-gated ion channel ELIC. Functional characterization demonstrates that nanobodies can act as positive or negative allosteric modulators. Crystal structures reveal that ELIC nanobodies can interact via distinct epitopes, including accessible parts of allosteric binding sites previously discovered in the extracellular ligand binding domain and bound by small molecules. One potentially attractive site for further development of allosteric modulators is the vestibule site, which can be targeted not only with nanobodies or small molecules in ELIC, but also by chemical modification of engineered cysteines as demonstrated in the human 5-HT_3A_ receptor. The vestibule site offers opportunities to further develop both positive and negative modulators, as well as to exploit subtype-specific access to certain receptors. These results pave the way for the future development of novel therapeutics that can modulate channel activity in pLGIC-related disorders. An attractive path is to expand the currently available repertoire of therapeutics with pharmacologically active nanobodies against human pLGICs.

## EXPERIMENTAL PROCEDURES

### Production of nanobodies against ELIC

A llama was immunized with 2 mg in total of purified wild type ELIC protein over a period of 6 weeks using a previously established protocol (Pardon et al. 2014). Briefly, from the anti-coagulated blood of the immunized llama, lymphocytes were used to prepare cDNA. This cDNA served as a template to amplify the open reading frames coding for the variable domains of the heavy-chain only antibodies, also called nanobodies. The PCR fragments were ligated into the pMESy4 phage display vector and transformed in *E. coli* TG1 cells. A nanobody library of 1.1×10^8^ transformants was obtained. For the discovery of ELIC specific nanobodies, wild type ELIC was solid phase coated directly on plates and phages were recovered by limited trypsinization. After two rounds of selection, periplasmic extracts were made and subjected to ELISA screens, seven different families were confirmed by sequence analysis. All clones were produced and purified as previously described (Pardon et al. 2014).

### Nanobody expression and purification

A series of nanobodies were individually expressed in the periplasm of *E. coli* strain WK6, which was grown in TB media supplemented with 0.1 mg/ml carbenicillin, 0.1% glucose and 2 mM MgCl_2_ to an absorbance A_600_∼0.7 at 37°C. The culture was induced with 1 mM isopropyl β-D-1-thiogalactopyranoside (IPTG) and incubated in an orbital shaker overnight at 28°C. Cells were harvested by centrifugation, resuspended in TES buffer (200 mM TRIS, pH 8.0; 0.5 mM EDTA; 500 mM sucrose supplemented with 40 mM imidazole) and incubated for 1 h. To this fraction 4 times diluted TES buffer was added and incubated for 1 h. This fraction was cleared by centrifugation at 10,000 g. The supernatant was incubated with Ni Sepharose 6 Fast Flow resin (GE Healthcare) and incubated for 1 h at room temperature. The beads were washed with buffer containing 20 mM TRIS pH 8.0, 300 mM NaCl and 40 mM imidazol. Protein was eluted with the same buffer supplemented with 300 mM imidazole. The eluted protein was concentrated to less than 1 ml on a 3 kDa cut-off Vivaspin concentrating column (Sartorius) and further purified on a Superdex 75 10/300 GL column (GE Healthcare) equilibrated with 10 mM Na-phosphate (pH 8.0) and 150 mM NaCl. Peak fractions corresponding to nanobody were pooled and spin-concentrated to ∼50 mg/ml.

### Automated voltage-clamp recordings of ELIC

For expression of ELIC in *Xenopus* oocytes we used the pGEM-HE expression plasmid (Liman, Tytgat, and Hess 1992) containing the signal sequence of the human α7 nAChR followed by the mature ELIC sequence, as previously described (Spurny et al. 2012). After plasmid linearization with *Nhe*I, capped mRNA was transcribed *in vitro* using the mMESSAGE mMACHINE T7 transcription kit (ThermoFisher). 2 ng of mRNA per oocyte was injected into the cytosol of stage V and VI oocytes using the Roboinject automated injection system (Multi Channel Systems). Oocyte preparations and injections were done using standard procedures (Knoflach, Hernandez, and Bertrand 2018). Injected oocytes were incubated in ND96-solution containing 96 mM NaCl, 2 mM KCl, 1.8 mM CaCl_2_, 2 mM MgCl_2_ and 5 mM HEPES, pH 7.4, supplemented with 50 mg/L gentamicin sulfate. One to five days after injection, electrophysiological recordings were performed at room temperature by automated two-electrode voltage clamp with the HiClamp apparatus (Multi Channel Systems). Cells were superfused with standard OR2 solution containing 82.5 mM NaCl, 2.5 mM KCl, 1.8 mM CaCl_2_, 1 mM MgCl_2_ and 5 mM HEPES buffered at pH 7.4. Cells were held at a fixed potential of −80 mV throughout the experiment. Agonist-evoked current responses were obtained by perfusing oocytes with a range of GABA concentrations in OR2 solution. Different nanobodies diluted into OR2 solution were tested at a range of concentrations by pre-incubation with nanobody alone and followed by a co-application of nanobody with 5 mM GABA. Data acquired with the HiClamp were analyzed using the manufacturer’s software (Multi Channel Systems). Concentration-activation curves were fitted with the empirical Hill equation as previously described (Spurny et al. 2012).

### ELIC purification and crystallization of ELIC-nanobody complexes

Purified ELIC protein was produced as previously described, but with minor modifications (Spurny et al. 2012). In brief, the ELIC expression plasmid was transformed into the C43 *E. coli* strain and cells were grown in LB medium. Protein expression was induced with 200 µM isopropyl β-D-1-thiogalactopyranoside (IPTG) and incubated in an orbital shaker at 20°C overnight. After cell lysis, membranes were collected by ultracentrifugation at 125,000 x g and solubilized with 2% (w/v) anagrade n-undecyl-β-D-maltoside (UDM, Anatrace) at 4°C overnight. The cleared supernatant containing the solubilized MBP-ELIC fusion protein was purified by affinity chromatography on amylose resin (New England Biolabs). Column-bound ELIC was cleaved off by 3CV protease in the presence of 1 mM EDTA + 1 mM DTT at 4°C overnight. A final purification step was carried out on a Superdex 200 Increase 10/300 GL column (GE Healthcare) equilibrated with buffer containing 10 mM Na-phosphate pH 8.0, 150 mM NaCl, and 0.15% n-undecyl-β-D-maltoside (UDM, Anatrace). Peak fractions containing pentameric ELIC were pooled, concentrated to ∼10 mg/mL and relipidated with 0.5 mg/mL *E. coli* lipids (Avanti Polar Lipids). Nanobodies were added at a 20% molar excess calculated for monomers and incubated at room temperature 2 h prior to setting up crystallization screens with a Mosquito liquid handling robot (TTP Labtech). Crystals for the ELIC complex with PAM-Nb grew at room temperature in the presence of 0.1 M GABA, 0.2 M Ca(OAc)_2_, 0.1 M MES buffer pH 6.5 and 10% PEG8000. Crystals for the ELIC complex with NAM-Nb grew at room temperature in the presence of 0.1 M Na_2_SO_4_, 0.1 M bis-trispropane pH 8.5 and 10% PEG3350. Crystals were harvested after adding cryo-protectant containing mother liquor gradually supplemented with up to 25% glycerol in 5% increments. Crystals were then plunged into liquid nitrogen and stored in a dewar for transport to the synchrotron.

### Structure determination of ELIC-nanobody complexes

Diffraction data for the ELIC+PAM-Nb structure were collected at the PROXIMA 1 beamline of the SOLEIL synchrotron (Gif-sur-Yvette, France). Diffraction data for the ELIC+NAM-Nb structure were collected at the X06A beamline of the Swiss Light Source (Villigen, Switzerland). Both structures were solved by molecular replacement with Phaser in the CCP4 suite (Winn et al. 2011) using the published structures for the ELIC pentamer (pdb 2vl0) and a GPCR nanobody (pdb 3p0g) as search templates. For the ELIC+PAM-Nb structure, the asymmetric unit contains one ELIC pentamer with 5 nanobodies bound (one to each subunit). For the ELIC+NAM-Nb structure, the asymmetric unit contains 2 ELIC pentamers with 1 nanobody bound to each pentamer. The electron density for one of these nanobody molecules is not well defined, suggesting partial occupancy at this pentamer, and therefore the atom occupancies for this nanobody were manually set to 40% during structure refinement. The data set for the ELIC+NAM-Nb was anisotropic with data extending to ∼3.15 Å in the best direction and ∼3.5 Å in the worst. To correct for anisotropy the unmerged reflections from XDS where uploaded to the STARANISO server (Paciorek et al. 2018) and automatically processed using CC1/2 > 30 and I/σ > 2 as resolution cut-off criteria. The merged and scaled data set from this procedure extends to a resolution of 3.25 Å and the statistics produced by the STARANSIO server are shown in the crystallographic table (supplementary Table 1). Structures were improved by iterative rounds of manual rebuilding in Coot (Emsley et al. 2010) and automated refinement in Buster (Smart et al. 2012) or Refmac (Winn et al. 2011). Structure validation was carried out in PDB-REDO (Joosten et al. 2014) and MolProbity (V. B. Chen et al. 2010). Figures were prepared with PyMOL (Schrödinger). Pore radius profiles were made using CHAP (Klesse, n.d.). Simulated annealing omit maps were calculated in PHENIX and are shown for the nanobody-ELIC interaction region (supplementary Figure S1-S2).

### Cysteine-scanning mutagenesis and voltage-clamp recordings of 5-HT_3A_R

Stage V-VI *Xenopus* oocytes were purchased from Ecocyte (Germany) and stored in ND-96 (96 mM NaCl, 2 mM KCl, 1.8 mM CaCl_2,_ 1 mM MgCl_2_, 5 mM HEPES, pH 7.5) containing 2.5 mM sodium pyruvate, 50 mM gentamicin and 0.7 mM theophylline. cDNA encoding human 5-HT_3A_R was cloned into the pGEM-HE expression plasmid (Liman, Tytgat, and Hess 1992). Mutants were engineered using the QuikChange mutagenesis kit (Agilent) and confirmed by sequencing. cRNA was *in vitro* transcribed from linearized pGEM-HE cDNA template using the mMessage mMachine T7 Transcription kit (ThermoFisher). Oocytes were injected with 50 nl of ∼400 ng/µl cRNA, and currents were recorded 18-48 hours post-injection. 5-HT_3A_R current recordings were obtained using a Roboocyte voltage-clamp system (Multi Channel systems) at a constant voltage clamp of −60 mV. Oocytes were perfused with ND-96 with no added calcium, and 5-HT (creatinine sulphate complex, Sigma) was diluted in this media for obtaining concentration-response curves. MTSEA-biotin (Biotium) was diluted immediately prior to application into calcium-free ND-96 solution at a concentration of 2 mM from a stock solution of 500 mM in DMSO. Analysis and curve fitting was performed using Prism v4.03 (GraphPad Software). Concentration-response data for each oocyte were normalized to the maximum current for that oocyte.

## ACKNOWLEDGMENTS

We thank beamline staff at the SOLEIL synchrotron and Swiss Light Source for assistance with data collection. SBO/IWT-project 1200261 and FWO-project G.0762.13 were awarded to JS and CU. Additional support was from KU Leuven OT/13/095, C32/16/035 and C14/17/093 to CU.

## TRANSPARENT REPORTING STATEMENTS

All *Xenopus* electrophysiology experiments were repeated 4-8 times. The number of “n” is mentioned in the relevant sections of the main text. We define each separate oocyte recording as a biological repeat. No data were excluded, unless the oocyte gave no detectable current. All electrophysiology experiments were conducted on automated devices, either the HiClamp or the Roboocyte, so essentially there was no human bias in recording of these data. Data are presented as the mean ± standard error of the mean (SEM) with the raw data points overlaid as a dot plot. Statistical comparison between groups of data was performed using an unpaired two-tailed *t* test and the significance value *P* is mentioned in the relevant sections of the manuscript.

The X-ray diffraction data sets were collected from single crystals and typically the data set with the highest resolution was used for structural elucidation. Equivalent reflection data were recorded multiple times in agreement with the rotational symmetry of the crystal packing. The relevant data multiplicity value for each data set is mentioned in the crystallographic table (supplementary info). All aspects of X-ray data collection, integration, scaling and merging were fully automated so human bias was excluded. No data were excluded.

## DATA AVAILABILITY

Atomic coordinates and structure factors have been deposited with the Protein Data Bank under accession numbers 6SSI for the ELIC+PAM-Nb structure and 6SSP for the ELIC+NAM-Nb structure. The raw X-ray diffraction images for both data sets have been deposited on datadryad.org under accession number doi:10.5061/dryad.pv4097s. A source file is submitted with the data plotted in Figure 1a-b, Figure 2c and Figure 4c-d-e.

## Supplementary information

**Table 1.** Crystallographic and refinement statistics.

|  | ELIC+PAM-Nb | ELIC+NAM-Nb |
| --- | --- | --- |
| Crystallographic statistics |  |  |
| Beamline | PROXIMA 1 (SOLEIL) | X06A (SLS) |
| Wavelength (Å) | 0.97857 | 0.9999 |
| Spacegroup | $P2_1$ | $P1$ |
| $a, b, c$ (Å) | 105.6, 146.56, 140.24 | 121.73, 122.29, 128.15 |
| $\alpha, \beta, \gamma$ (°) | 90, 111.39, 90 | 71.02, 64.20, 61.66 |
| Resolution limits (Å) | 49.67 - 2.59 (2.65 - 2.59) | 48.32 - 3.25 (3.34 - 3.25) |
| $R_{merge}$ (%) | 6.1 (122.1) | 5.2 (95.4) |
| $R_{meas}$ (%) | 7.8 (157.1) | 7.4 (134.9) |
| $R_{pim}$ (%) | 4.9 (97.8) | 5.2 (95.3) |
| $\langle I/\sigma \rangle$ | 13.4 (1.0) | 5.5 (0.5) |
| $CC_{1/2}$ (%) | 99.9 (45.8) | 99.7 (42.9) |
| Multiplicity | 4.8 (4.6) | 1.8 (1.6) |
| Completeness (%) | 99.7 (95.2) | 95.1 (89.4) |
| Total number of reflections | 594559 (26957) | 147646 (6664) |
| Number unique reflections | 123770 (5799) | 83183 (4160) |
| Refinement and model statistics |  |  |
| $R_{work}$ (%) | 22.97 | 24.80 |
| $R_{free}$ (%) | 24.59 | 26.43 |
| Rmsd bond distance (Å) | 0.008 | 0.008 |
| Rmsd bond angle (°) | 0.94 | 0.97 |
| Ramachandran analysis |  |  |
| Outliers (%) | 0.05 | 0.16 |
| Favored (%) | 97.16 | 96.53 |
| Poor rotamers (%) | 0.63 | 0.77 |
| Molprobability score | 1.54 (100rd percentile) | 1.78 (100rd percentile) |

**Supplementary Figure S1.**
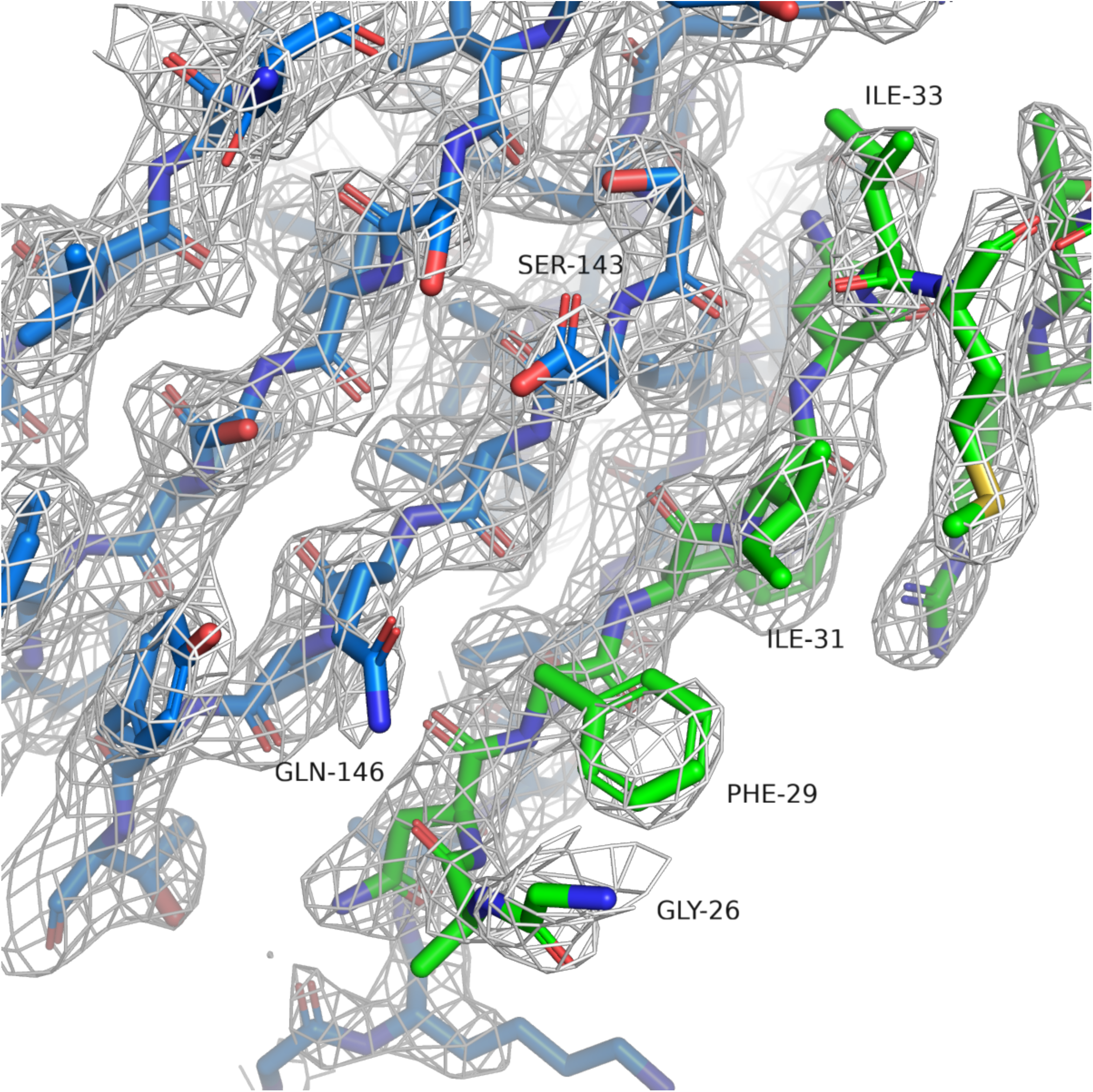
Simulated annealing omit map for ELIC+PAM-Nb structure. The white mesh shows a simulated annealing omit map calculated in PHENIX and contoured at a sigma level of 1.0 around a 12 Å sphere zoom of the ELIC-Nb interface. In the omit calculation an omit box around the entire chain F (green) of the PAM-Nb was selected. ELIC is shown in blue and selected residues of the interaction interface are indicated.

**upplementary Figure S2.**
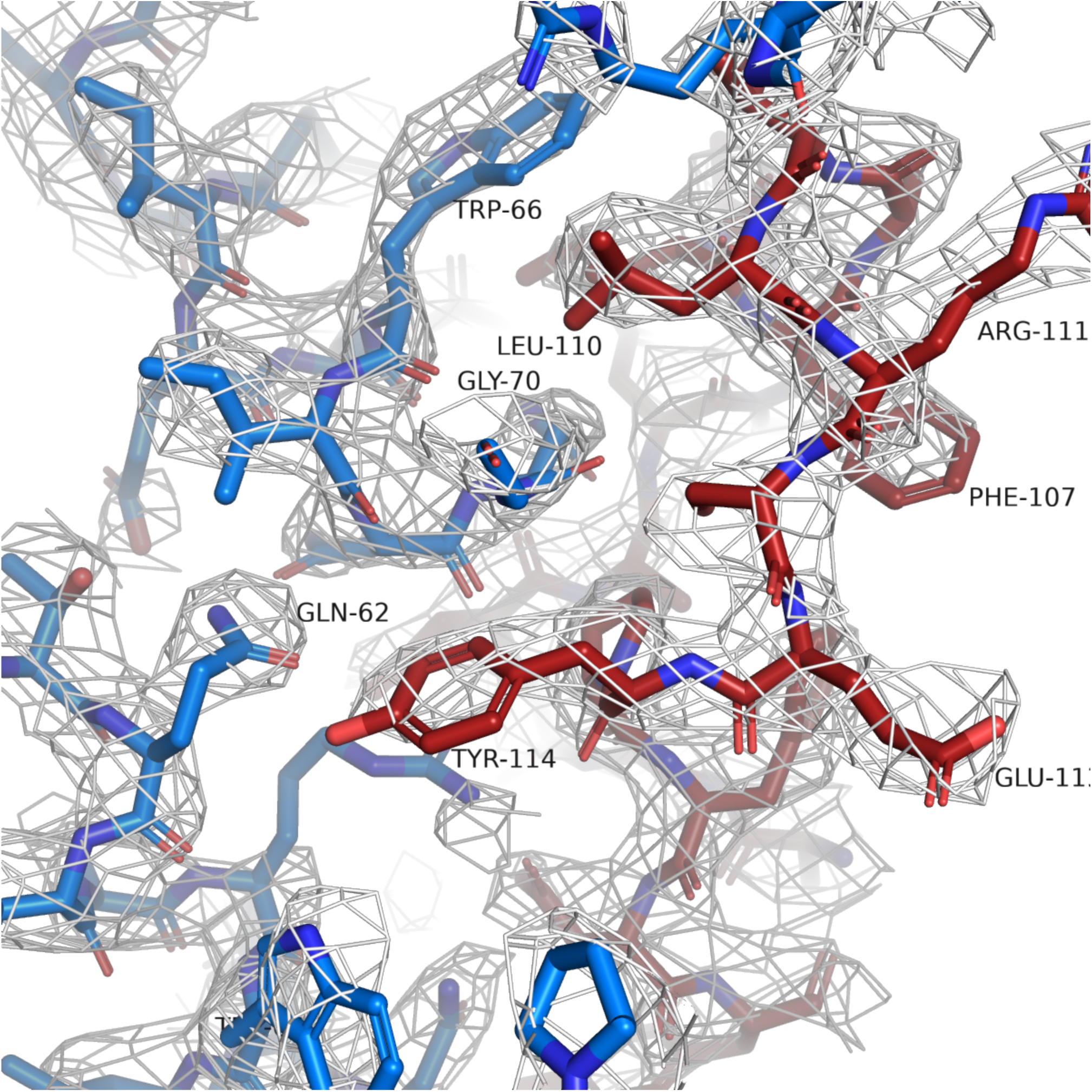
Simulated annealing omit map for ELIC+NAM-Nb structure. The white mesh shows a simulated annealing omit map calculated in PHENIX and contoured at a sigma level of 1.0 around a 12 Å sphere zoom of the ELIC-Nb interface. In the omit calculation an omit box around the entire chain K (red) of the PAM-Nb was selected. ELIC is shown in blue and selected residues of the interaction interface are indicated.

